# Structure and physiological investigation of arginylated actin

**DOI:** 10.1101/2024.06.12.598685

**Authors:** Clyde Savio Pinto, Saskia E. Bakker, Andrejus Suchenko, Hamdi Hussain, Tomoyuki Hatano, Karuna Sampath, Krishna Chinthalapudi, Masanori Mishima, Mohan Balasubramanian

## Abstract

Actin is an evolutionarily conserved cytoskeletal protein with crucial roles in cell polarity, division, migration, and muscle contraction. Actin function is regulated in part by posttranslational modifications. One such modification in non-muscle cells is arginylation, in which an arginine residue is added to the N-terminus of β-actin. What is the structure of arginylated β-actin (R-β-actin), are its interactions with other proteins altered and what phenotypes result when R-β-actin is the sole actin isoform present in the cell? Here we report the 4.2 Å structure of ADP-bound human R-β-actin filaments, the overall structure of which is nearly identical to the filaments made of non-arginylated actin. *In vitro* functional assays using isoform-pure actins with defined post-translational modifications reveal that the interaction between myosin-II and actin is altered upon actin arginylation, due to frequent detachment of myosin-II from R-actin filaments. *In vivo*, we find that replacement of the only actin gene in *Schizosaccharomyces pombe* with a synthetic gene encoding R-Sp-actin reduces Arp2/3-based actin patches while thickening the formin-induced actin. Furthermore, consistent with altered interactions between myosin-II and R-actin filaments, the assembly and constriction of cytokinetic actomyosin ring are perturbed in the R-Sp-actin cells. Thus, despite the overall structural similarity of arginylated and non-arginylated actin filaments, actin arginylation affects actin filament assortment into distinct subcellular structures and its interaction with myosin II.

## Introduction

Actin is one of the most conserved and abundant eukaryotic cytoskeletal proteins and plays a key role in major cellular functions such as motility, division, polarity, and muscle contraction (***Dominguez and Holmes***, 2011; ***Holmes*** 1998; ***Kabsch and Holmes***, 1995; ***Pollard and Cooper***, 2009). The structure of actin has been determined by cryogenic electron microscopy revealing near-atomic resolution actin filament structures (***Arora et al***., 2023; ***Avery et al***., 2017; ***Belyy et al***., 2020; ***Chou and Pollard***, 2019; ***Kumari et al***., 2020; ***Raunser***, 2017; ***von der Ecken et al***., 2016)), which have built on beautiful work elucidating the structure of actin monomers using X-ray crystallography (***Holmes et al***., 1990; ***Kabsch et al***., 1990; ***Schutt et al***., 1993). Actin organizes into two helical strands, which undergo continuous polymerization and depolymerization. Actin isoforms have posttranslational modifications (PTM) that appear to facilitate specific cellular functions (***Terman and Kashina***, 2013; ***Varland et al***., 2019). Amongst these modifications for the human non-muscle β-actin are the N-terminal acetylation and His-73 methylation (***Terman and Kashina***, 2013; ***Varland et al***., 2019). In addition, a subset of β-actin undergoes a process termed arginylation in which an arginine residue is added to the N-terminus (***Karakozova et al***., 2006b; ***Kashina***, 2014; ***Lian et al***., 2014; ***Terman and Kashina***, 2013). It is now clear that His-73 methylation, carried out by the SETD3 family of methyl transferases regulates Pi release following ATP hydrolysis within the filament (***Dai et al***., 2019; ***Hintzen et al***., 2021; ***Kwiatkowski et al***., 2018; ***Lappalainen***, 2019; ***Wilkinson et al***., 2019). N-terminal acetylation of actin, mediated by the NAA80 enzyme, plays an important role in actin polymerization and depolymerization kinetics and affects cellular structures such as filipodia and lamellipodia (***Aksnes et al***., 2018; ***Arnesen et al***., 2018; ***Beigl et al***., 2020; ***Drazic et al***., 2018, 2022; ***Ree et al***., 2020). β-actin arginylation, which is achieved by the arginyl-tRNA-protein transferase Ate1 following the removal of the N-terminal acetylated aspartic acid residue, has been proposed to regulate lamellipodia formation, cell size, and cell spreading, but the exact biochemical and structural mechanisms are unknown (***Karakozova et al***., 2006b; ***Saha et al***., 2010a).

In this study, to address the structural and physiological mechanisms impacted by actin arginylation, we used singleparticle cryo-electron microscopy combined with *in vitro* analysis of the actin-myosin interaction as well as *in vivo* analysis in the genetically tractable fission yeast *Schizosaccharomyces pombe*. Our findings identify differences between arginylated and non-arginylated forms in actin filament organisation and interactions of the filaments with myosin.

## Results

### No drastic effect of arginylation on actin and actin filament structures

Using a method that we recently developed, we expressed and purified from *Pichia pastoris* large amounts of recombinant human R-β-actin (***Hatano et al***., 2018) in which His73 was methylated by co-expression of human SETD3 (***Hatano et al***., 2020). To assess the effect of N-terminal argynilation on the actin filament, we solved the structure of fully argynilated actin filaments by cryo-EM. Globular R-β-actin was polymerized, plunge-frozen, and screened to evaluate filament formation and ice thickness. Following optimization of filament assembly conditions, we collected 2285 images where the actin filaments are clearly identifiable in the raw micrographs (Fig. 1. A). Individual filament images were boxed into overlapping segments and 50 classes were generated. The wide and narrow projections of the filaments are resolved in the class averages (Fig. 1. B). Following this, using the helical reconstruction approach (***He and Scheres***, 2017), we obtained a 3D map of the ADP-bound human R-β-actin to a resolution of 3.5 Å for the subunit. In this map, displayed with the pointed end of the filament positioned to the top, we observed well-resolved secondary structure elements. Local resolutions range from 2.2 Å for the core region to 4.2 Å for the flexible, solvent-exposed areas including the D-loop and N-terminus. The refined map shows a helical twist of 166.8 degrees and a rise of 28.1 Å consistent with values reported previously (Fig. 1. C) (***Arora et al***., 2023; ***Chou and Pollard***, 2019; ***Galkin et al***., 2015).

**Figure 1.**
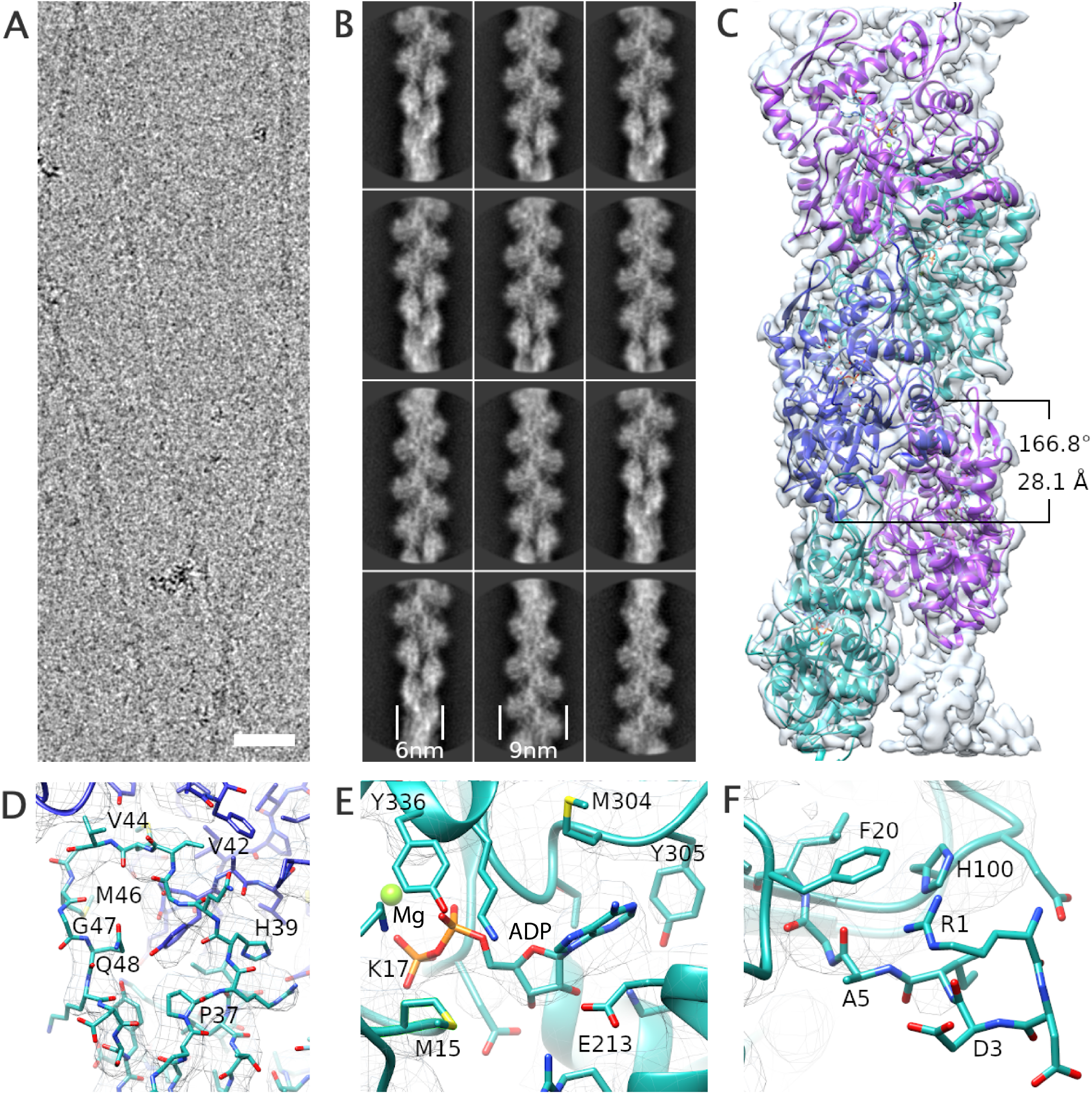
The structure of R-β-actin. (A) Representative cryo-TEM image of human filamentous R-β-actin. Scale bar 20 nm. (B) The 2D Class averages obtained from 50 individual regions selected from the filaments, showing the wide and narrow projections of filaments (indicated with lines). (C) Three-dimensional reconstruction of human F-actin showing 5 actin subunits in different colours. (D) The DNAse I-binding loop (D-loop) of R-β-actin shows the fit of the protein backbone in the density. Residues in this region are shown in stick representation. (E) Ribbon diagram of the filamentous R-β-actin ADP-binding site, with ADP and interacting residues shown in stick representation. (F) The N-terminus region of R-β-actin from residue Ile-4 is shown. Electron density fit to the residues 1 to 3 is missing as this region is not resolved.

An atomic model was built into the density map using a rigid-body fit of the published chicken skeletal muscle actin (***Galkin et al***., 2015) as a starting point. At the present resolution, we were able to reliably trace the backbone (Fig. 1. C) and resolve the density of the side chains of the larger residues (Fig. 1. D) and Mg^2+^-ADP (Fig. 1. E). Compared to the core of the subunit, we observe a weaker backbone density for residues Met46 and Gly47 in the DNase I-binding loop (D-loop) (Fig. 1. D). The 2 N-terminal residues are not resolved in the electron density (Fig. 1. F) indicating the N-terminus is flexible. This is consistent with the lower resolution of this region as shown by the local resolution analysis.

The structure of the arginylated human F-actin is consistent with previously published F-actin structures, including the recently published human beta-actin structure (***Arora et al***., 2023) (Fig. 2). The root-mean-square deviation (RMSD) between R-β-actin and human Ac-β-actin (8DNH) is 0.851 Å (Fig. 2. A) and the active site is consistent with previous structures (Fig. 2. B). A comparison of the N-terminal residues shows the large number of negatively charged residues in β-actin, which are reduced by the arginylation (see also Fig. 2. C). We then compared R-β-actin with previously published chicken and rabbit skeletal muscle F-actin (Fig. S1. A-C) (***Chou and Pollard***, 2019; ***Galkin et al***., 2015). This shows a backbone RMSD of 0.965 Å between R-β-actin filament structure determined in this work and 6DJO (chicken, (***Chou and Pollard***, 2019)) and 1.248 Å between the structure of R-β-actin and 3J8I (rabbit, (***Galkin et al***., 2015)). Despite the higher sequence similarity between the rabbit and chicken actin models, the RMSD between the two at 1.353 Å is lower than between either of those structures and our model of human F-actin.

**Figure 2.**
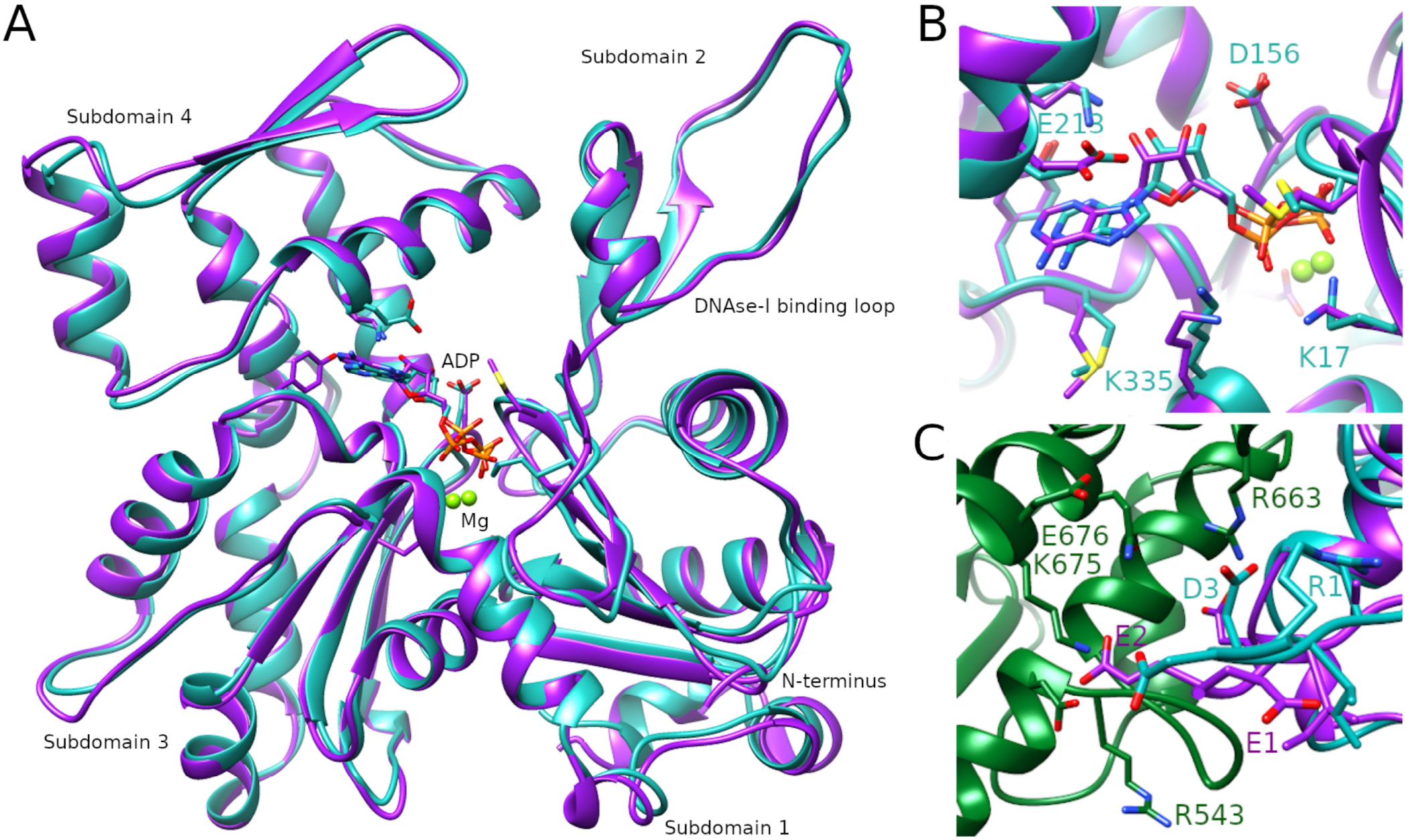
Comparison of the R-β-actin and Ac-β-actin structures and their interaction with myosin II. (A) Ribbon diagram comparing the structure of R-β-actin presented here (cyan) with the previously published structure of human Ac-β-actin (magenta). (B) Close up view comparing the structure of the active site of R-β-actin with previously published structures of human filamentous Ac-β-actin. (C) Comparison of free R-β-actin (cyan) and human filamentous Ac-β-actin (magenta) bound to myosin (green), showing the interaction of the N-terminal residues with the myosin.

### Myosin-II frequently detaches from R-actin filaments during motility

Actin arginylation following removal of the N-terminal acetylated aspartate increases the positive electrostatic surface charge by 3 units. In the actomyosin complex, the N-terminal tail of actin is closely located to the positively charged surface patches on myosin (Fig. 3. A). The N-terminal residues of actin have been reported to be involved in the functional/physical interaction with myosin (***Arora et al***., 2023; ***Behrmann et al***., 2012; ***von der Ecken et al***., 2016). Thus, we then examined the influence of actin arginylation on actin-myosin interaction using *in vitro* assays with purified proteins.

**Figure 3.**
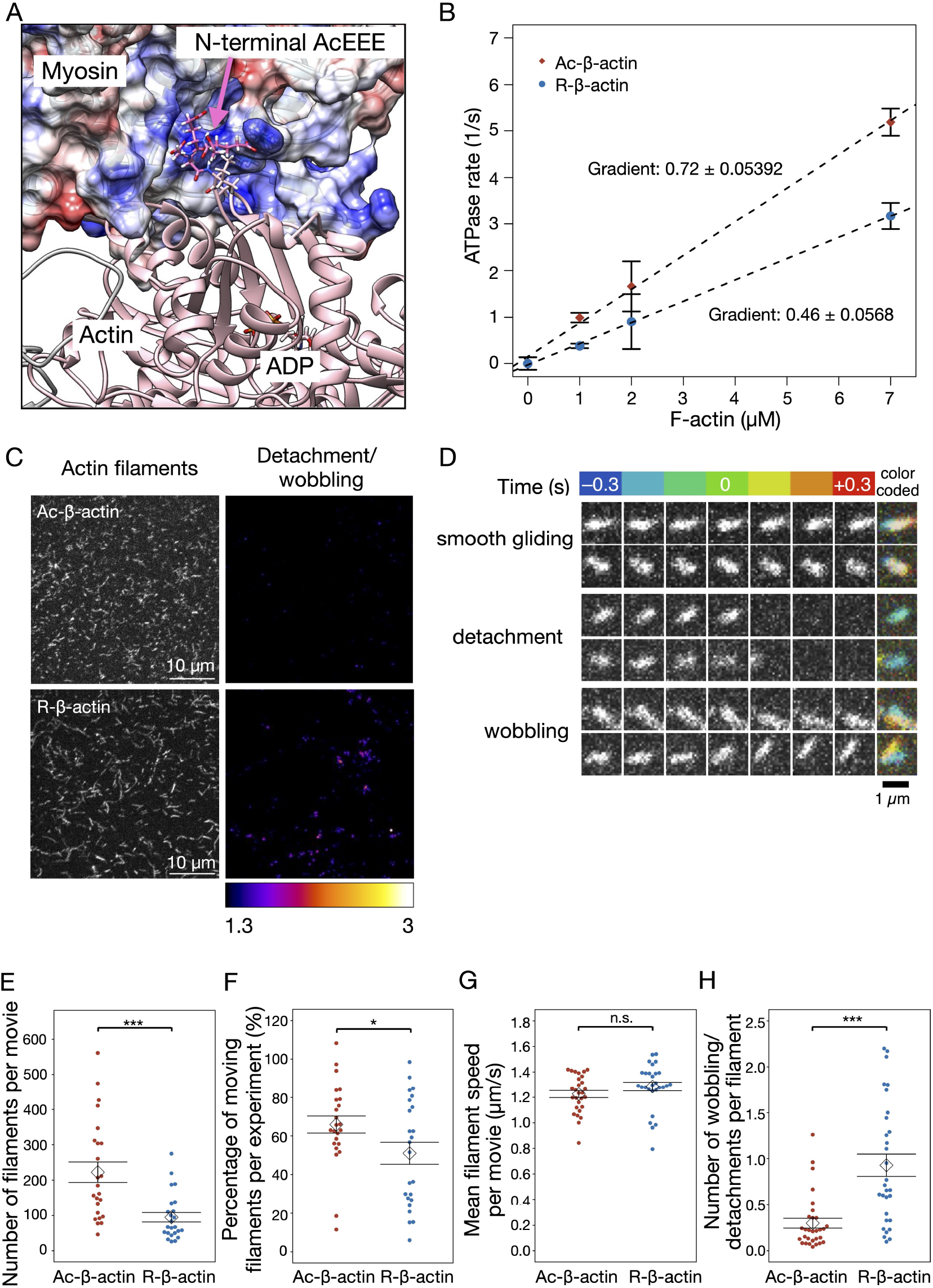
The stability of the interaction between myosin and actin filaments is weakened by arginylation of actin. (A) A model of the actomyosin complex, indicating the electrostatic interaction between the negatively charged actin N-terminal tail and the positively charged surface of myosin. A snapshot from a molecular dynamic simulation based on a cryo-EM structure (PDB:5JLH (***von der Ecken et al***., 2016)) is displayed with the N-terminal tail of gamma-actin (AcEEEI) in magenta and the surface Coulomic potential of myosin IIc in a red (0 to –10 kcal/mol) and blue (0 to +10 kcal/mol) linear scale. (B) The steady-state rate of ATP hydrolysis by cardiac myosin measured in the presence of Ac- or R-actin filaments. (C) The surface motility assay with immobilized cardiac myosin and Alexa Fluor-488-labelled filaments of Ac- or R-actin. The left panels show the total filaments landed in the presence of 1 mM ATP. The right panels indicate the detachment or wobbling of filaments. The ratios between the pixel intensities of the n-th and (n+1)-th images of the movie were then calculated. These ratio images were maximum-projected through the time points and presented on a linear scale from 1.3 to 3 using the pseudocolor ‘FIRE’ lookup table. (D) Examples of the detachment and wobbling of actin filaments. Indicated time frames before and after the detected event were shown as well as the maximum projection with temporal color coding (blue to red). (E-H) The quantification of the myosin-driven motility of Ac- or R-actin filaments. Each dot corresponds to experiment or a movie of 20 s duration. Mean (diamond) and the standard error (error bars) are indicated. (E) Number of filaments per movies. (F) Percentage of the moving filaments. (G) Average speeds of the moving filaments per movie. (H) Frequency of detachment or wobbling per filament.

First, we assessed if R-β-actin assembled filaments with equal efficiency compared to Ac-β-actin, as determined using a pelleting assay. No significant difference was detected between R-β-actin and Ac-β-actin after polymerization in the fraction of filaments sedimented following centrifugation (Figure. S2. A). When loaded onto myosin-II coated coverslips in the absence of ATP, the number of filaments attached, and their length distributions were similar between the R-β-actin and Ac-β-actin filaments (Fig. S2. B,C,D). These observations indicate that the effect of arginylation on the *in vitro* actin polymerization is minimal.

Interaction with actin activates the ATPase domain of myosin leading to hydrolysis of ATP, which produces energy for mechanical work. We assessed the actin-activated ATPase activity of cardiac myosin-II by measuring the increase of free phosphate using a spectrophotometric assay. Although both polymerized Ac-β-actin and R-β-actin promoted myosin ATPase activity in concentration-dependent manners, the degree of stimulation by R-β-actin was about 50% that of Ac-β-actin (Fig. 3. B). This indicates that R-β-actin is less effective in activating the ATPase activity of myosin-II compared to Ac-β-actin.

Myosin is an actin-based motor protein whose power stroke translocates actin filaments when the myosin is anchored. We investigated the influence of the arginylation on myosin motor activity by observing the motility of fluorescently labelled R-β-actin (95% R-actin + 5% fluorescently labelled Ac-actin) and Ac-β-actin (100% acetylated of which 5% was fluorescently labelled) filaments on the coverslip surface coated with cardiac myosin (Video 1). Although, as described above, the initial surface landing of R-actin and Ac-actin filaments in the absence of ATP was similar, they behaved differently upon the addition of ATP. First, a higher number of Ac-β-actin filaments were retained on myosin-II-coated coverslips compared to R-β-actin filaments following ATP addition (Fig. 3. C,E). Consequently, the number of motile actin filaments was reduced in the R-β-actin sample compared to the Ac-β-actin sample (Fig. 3. F). The R-β-actin filaments retained on the surface exhibited motility at a comparable velocity to the Ac-β-actin filaments (Fig. 3. G). However, consistent with the reduced surface retention, the R-β-actin filaments showed abrupt displacement (‘wobbling’) or detachment from the surface much more frequently than the Ac-β-actin filaments (Fig. 3. C,D,H). Collectively, these observations suggest that the negative charge in the N-terminal actin tail, which is reduced by the arginylation, contributes to persistent and productive interaction between F-actin and myosin-II, preventing unfavourable detachment of actin filaments from myosin-II.

### Cells that have R-Actin as the sole actin are viable but show an altered actin distribution and actomyosin ring function

The structural and biochemical work strongly suggested that actin arginylation compromises actin-myosin-II interaction. We designed the *in vivo* experimental setup that allows us to assess the physiological consequences of replacing the entire cellular pool of actin with R-actin. Analysis of the function of actin modifications are complicated in mammalian cells due to the multiplicity of actin isoforms and modifications. To generate cells that expressed only R-actin we used the fission yeast *Schizosaccharomyces pombe. S. pombe* has only one actin (*act1*) which does not undergo N-terminal maturation. It also has both Arp2/3 and formin nucleated actin structures making it an ideal system in which to study the effect of actin arginylation on the actin cytoskeletal organization and function. To express R-actin in fission yeast cells, we used our established strategy to express and purify R-actin from methylotrophic yeast, *Pichia pastoris* (***Hatano et al***., 2018). Towards this goal, we engineered the endogenous *act1* locus of *S. pombe* so that it expresses a fusion protein of a Ubi4 tag and either wild-type actin or R-actin. The Ubi4 tag is cleaved off inside the cell resulting in either Wt-Act1 or R-Act1, *S. pombe* actin with the N-terminal tail starting with MEEE and REE, respectively. We modified the *act1* locus in the wild-type background (*wt*) as well as in the strains expressing rlc1-3GFP and mCherry-atb2 (strain referred to as ‘*rlc1-GFP*’, hereafter) as markers of the actomyosin ring and the mitotic spindle respectively (Fig. S3). The successful gene editing was confirmed by PCR and sequencing (Fig. S4).

Efficient removal of the Ubi4 tag (8.7 kDa) in *S. pombe* cells was confirmed by western blotting, which showed only a single band at the expected size for actin (42 kDa) (Fig. S5. A) in all the MEEE Act1 strains constructed. REE Act1 also appeared as a single band with a slightly higher mobility as shown previously with human R-β-actin purified from *P. pastoris* (***Hatano et al***., 2018). The N-terminal sequence of MEEE-Act1 and REE-Act1 in respective strains was also confirmed by mass spectroscopy (Fig. S5. B,C,D).

There was no obvious difference in the viability or growth rate of REE and MEEE strains in spot assays (Fig. S5. E). However, phalloidin staining in the wt background revealed a decrease in the area occupied by actin patches in the REE strain as opposed to the MEEE strain (Fig. 4. C,D). On the other hand, we qualitatively observed thicker cables in the REE strain (Fig. 4. C). Furthermore, time-lapse observation revealed significant differences between them in the timings of the cytokinesis stages and the morphologies of the actin cytoskeleton (Fig. 4). The cytokinetic actin ring (CAR) in fission yeast is assembled from acto-myosin nodes, which first appear and then coalesce into a ring (node coalescence phase), which then dwells without constricting for a time (dwell phase) and then constricts (ring constriction phase). Using myosin regulatory light chain fused to 3 GFPs (Rlc1-3GFP) as a marker for the CAR we observed that, in the REE strain, nodes took longer to coalesce into a CAR, and the CAR dwelled for a shorter period than in the MEEE strain (Fig. 4. A,B, Video 2). The ring constriction took longer in the REE strain, reflecting the reduced average rate of constriction (ring diameter divided by the time for constriction). Overall, the total time taken for cytokinesis from node appearance to the end of CAR constriction was increased in the REE strain. These observations, especially the slower ring constriction, suggest that myosin II function in the REE strain is altered.

**Figure 4.**
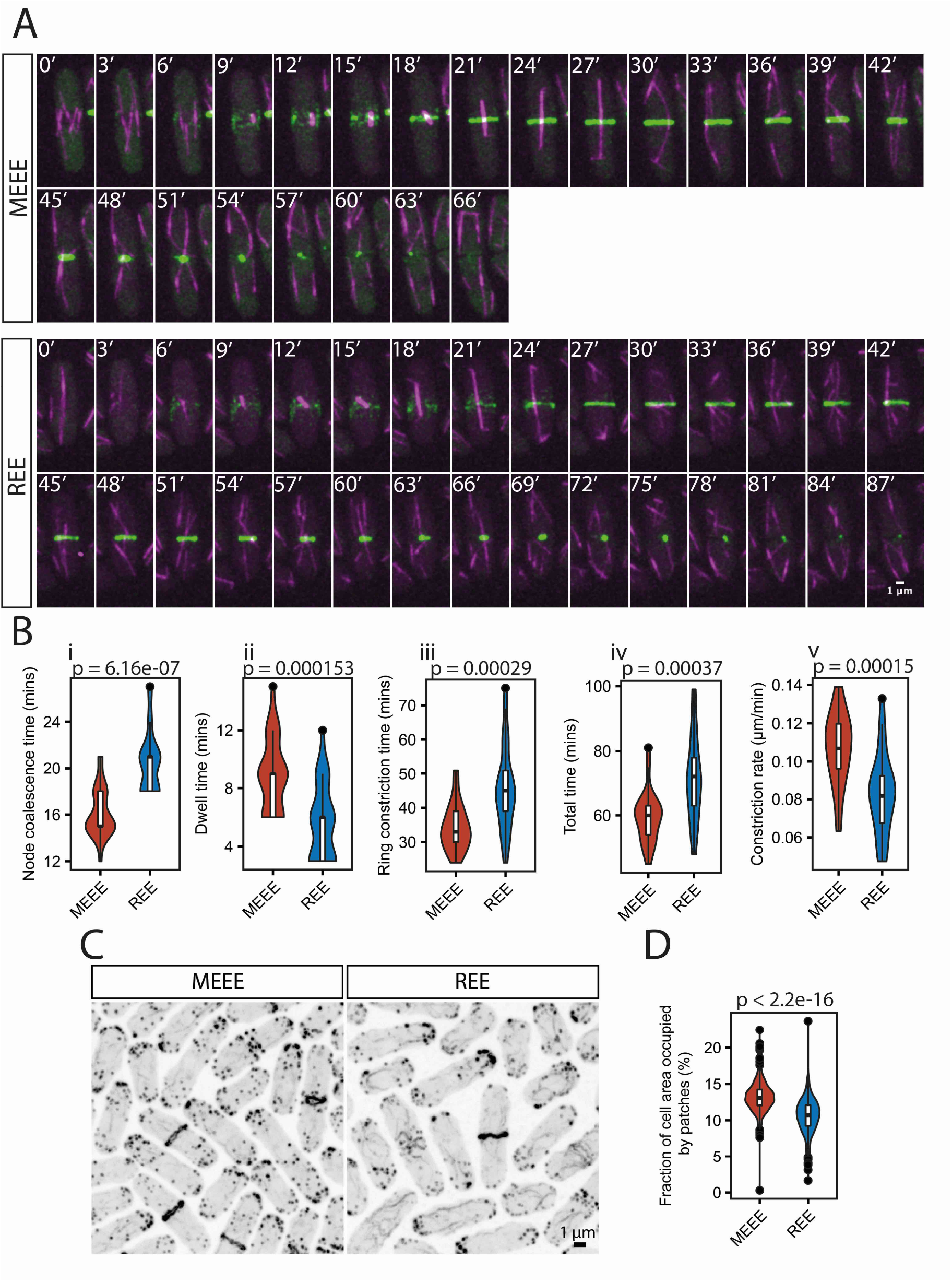
Cells with R-Actin as the sole actin have defects in actin distribution and cytokinesis. (A) Representative montages of time-lapse movies of live cells from the indicated strains in the *rlc1-GFP* background showing mCherry-Atb2 (magenta) and Rlc1-3GFP (green). Images were captured at 3 min intervals. (B) Violin plots of the analysis of ring constriction (n = 25 cells each) for (i) node coalescence (REE: mean = 20.52 mins; SD = 2.69; MEE: mean = 16.20; SD = 2.12; Wilcoxon test p = 6.16e-07), (ii) dwell time (REE: mean = 5.64 mins; SD = 2.64; MEEE: mean = 8.88; SD = 2.52; Wilcoxon test p = 0.000153), (iii) ring constriction time (REE: mean = 45.48 mins; SD = 12.27; MEEE: mean = 34.32 mins; SD = 6.66; T test p = 0.00029), (iv) total time (REE: mean = 71.64 mins; SD = 13.54; MEEE: mean = 59.40 mins; SD = 7.94; T test p = 0.00037) and (v) constriction rate (REE: mean = 0.083 μm/min; SD = 0.021; MEEE: mean = 0.106 μm/min; SD = 0.019; T test p = 0.00015). (C) Phalloidin stainings of the indicated strains in the *wt* background. (D) Violin plots of the area occupied by patches as a fraction of cell area for the two indicated strains in the *wt* background (REE: n = 425 cells; mean = 10.59%; SD = 2.35; MEEE: n = 462 cells; mean = 13.42%; SD = 2.03; Wilcoxon test: p < 2.2e-16). Scale bars in A, C are 1 μm.

## Discussion

In this study, we have resolved the structure of R-_β_-actin, discovered that arginylation of actin alters its interaction with myosin II and developed a fission yeast system in which arginylated actin is the sole genetically provided actin. Our comparison of the R-β-actin structure with those from our previous work with Ac-β-actin as well as other previously published actin structures, was not able to detect major differences in their overall structures. However, within our R-_β_-actin structure we were unable to resolve the N-terminus. This suggests that the N-termini of different subunits within the filament are in different conformations, indicating a higher flexibility in the N-terminal region. Despite the reduction in negative charge at the N-terminus compared to acetylated actin, it appears the N-terminal arginylation does not help to stabilise the N-terminus, at least in the ADP-bound structure. Like the previously published ADP bound F-actin structures, our work shows a less well-defined density in the D-loop where adjacent subunits interact (Fig. 1. D) (***Chou and Pollard***, 2019; ***Merino et al***., 2018). Chou and Pollard have shown that in chicken actin this happens specifically in the ADP bound structure suggesting that the flexibility increases after hydrolysis of the ATP during polymerization (***Chou and Pollard***, 2019). Merino et al showed a similar drop in resolution for the ADP-bound structure, which could be prevented in actin stabilised with jasplakinolide (***Merino et al***., 2018).

Previous work has shown functional differences between arginylated actin and acetylated actin in terms of polymerization and its functional interactions with actin binding proteins such as the Arp2/3 complex (*Chin et al*., 2022). Since, the structure that we obtained for R-_β_-actin cannot account for these differences though we were unable to sufficiently resolve the N-terminus and the D-loop, we suggest that the charge difference on the N-terminus in R-_β_-actin might be the cause of these differential interactions. Previous work has demonstrated that the quantum of negative charge on the N-terminus is important for force generation through the interactions of the negatively charged actin N-terminus with the positively charged myosin II (***Arora et al***., 2023; ***Behrmann et al***., 2012; ***Lu et al***., 2005; ***Miller et al***., 1996; ***von der Ecken et al***., 2016). Indeed, our *in vitro* experiments show that R-β-actin filaments frequently detach from myosin-II and actin activated myosin ATPase activity is reduced (potentially due to the untimely detachment of actin filaments from myosin-II). It is likely that the positively charged arginine decreases the affinity of myosin for the otherwise negatively charged actin N-terminus resulting in increased detachment of actin from myosin, and consequently decreased actin activated myosin ATPase activity and a reduced number of filaments showing gliding motility.

Most previous work that aims to study the role of R-actin in cells relies on knock-out or inhibition of the arginyl transferase enzyme *ate1* which is responsible for arginylating actin (***Karakozova et al***., 2006a; ***Saha et al***., 2012, 2010b). A potential limitation of this approach is that ATE1 is responsible for the arginylation of a broad range of molecules and thus it is difficult to know whether the observed phenotype is due to the loss of R-actin or the loss of other arginylation. Our results demonstrate the functional difference between Ac-actin and R-actin in the cell. One question that has not been resolved to date is if R-actin can perform all the functions of wt actin in cells or if it can only interact with a subset of actin binding proteins. Since actin’s interactions with the Arp2/3 complex, formins, and myosin are critical to cell function it is reasonable to assume that significantly compromised functional interactions with any of these proteins and actin will be deleterious or even lethal to cells. Interestingly, we found that cells were capable of survival with subtle actin patch, cable, and cytokinesis defects when REE-actin was the sole genetically provided actin in the cell. Actin patches are nucleated by the Arp2/3 complex (***Kaksonen et al***., 2003) suggesting that Arp2/3 based actin nucleation is likely compromised in the REE strains. This is consistent with defective Arp2/3 complex based actin nucleation of R-_β_-actin which we observed previously (***Chin et al***., 2022). Thicker cables are an indicator of failed myosin function or an anti-correlation between patches and cables in fission yeast (***Balasubramanian et al***., 1996, 1998; ***Burke et al***., 2014; ***Zambon et al***., 2020). It is very likely based on our *in vitro* analysis that the charge differences in REE-actin and its consequent decrease in myosin II interaction led to the cytokinesis defects that we observed in the REE strain. Based on this work we show that arginylation doesn’t alter the overall structure of actin. Nonetheless, we suggest that the change in charge induced on the N-terminus alters its local molecular scale interactions that determine its localisation and function.

## Materials and Methods

### Yeast strains, media and culture conditions

Fission yeast S. pombe strains used in this study have been listed at the end of the manuscript. Cells used for live imaging or staining were grown in YEA medium at 30 °C with shaking at 200 rpm. The MEEE and REE strains were generated using the SpEDIT system (***Torres-Garcia et al***., 2020). Briefly, an sg-RNA (Designed using CHOP-CHOP (***Labun et al***., 2019)) loaded pLSB plasmid was generated using the following primers in a goldengate cloning reaction (F:CTAGAGGTCTCGGACTACCCTCAAAAGACAAGACCAGTTTCGAGACCCTTCC R:GGAAGGGTCTCGAAACTGGTCTTGTCTTTTGAGGGTAGTCCGAGACCTCTAG)

Targeting the 5’UTR near the ATG site of act1. A homologous recombination template (HDR-template) containing the ubi4-MEEE-act1 (till the stop codon) or ubi4-REE-act1(till the stop codon) was generated using pcr from plasmids containing these sequences using the following primers and Phusion polymerase (New England Biolabs: M0530S) (F:AATCAACGGCTTCATACCACCTCAGCCAGCCGTGTTATAACTTACCGTTTACCAACTACATTTTTTGTAACGAACCAAAAAACCCTCAAAAGACAAGACCATGCAGATTTTCGTCAAGAC R: TTAGAAGCACTTACGGTAAACG)

The products were pcr purified using a Qiagen pcr purification kit (28104). The HDR-template and sgRNA-pLSB plasmid were transformed into the wt and rlc1-GFP strains and colonies were selected on YEA plates containing NAT antibiotic at 24 °C. The colonies were then transferred to YEA plates and screened using pcr to detect Ubi4 incorporation using the following genotyping primers (F:GCATTCTGCCGTGAAGTG R: GCTCAAAGTCCAAAGCGAC).

They were then sequenced using the service from GATC and the sequences were aligned using SnapGene® software (from Dotmatics; available at snapgene.com) and confirmed. Schematics related to DNA sequences were also made in Snapgene.

### Imaging of *S. pombe* strains

#### Spinning disk confocal microscopy

All imaging was performed using the Andor Revolution XD spinning disk confocal microscope. Imaging was performed at 30°C for all live imaging experiments. The spinning-disk confocal system consisted of a Nikon ECLIPSE Ti inverted microscope with a Nikon Plan Apo Lambda 100 x/1.45 NA oil immersion objective lens, a spinning-disk system (CSU-X1; Yokogawa), and an Andor iXon Ultra EMCCD camera. A a pixel size of 80 nm/pixel was used for acquiring the images using the Andor IQ3 software. Lasers with wavelengths of 488 nm or 561 nm were used to excite the fluorophores. The time-lapse images were acquired with a Z-step of 500 nm.

#### Preparation of cells for live imaging

Cells used for live imaging were grown in YEA medium at 30 °C with shaking at 200 rpm and grown to mid-log phase. 1 ml of cells were centrifuged at 450xg for 2 mins to concentrate them to 20-100 μl and mounted sandwiched between an agarose pad and a coverslip which was sealed with valap (a mixture of vaseline, lanolin and paraffin) and imaged. Images were collected at 3 min intervals.

#### Phalloidin staining of S. pombe strains

For phalloidin stainings, the cells were grown in YEA medium at 30 °C with shaking at 200 rpm. They were grown to mid-log phase. 50 ml of the culture was spun down at 3000 rpm for 1 min, washed once with PBS and spun again. 500 μl of PBS was added to the pellet and 500 ul of the resultant solution was taken for fixing. 500 μl of 8% PFA was added to the 500 μl of cells and mixed immediately and then shaken on a nutator for 45 mins. The cells were spun down at 845g for 1 min and the PFA was discarded. The cells were washed once with PBS and then kept in 1 ml PBS at 4 °C till processing.100 μl of these cells were taken and 1 ml of 1% PBT was added to them for 30 mins at RT. The cells were washed once with PBS and the pellet was resuspended in 10 μl 1:10 Phalloidin, 1 μg/ml DAPI in PBS and mounted between an agarose pad and coverslip for imaging.

#### Analysis of S. pombe images

Live images were analyzed manually in FIJI by CP for ringrelated parameters (***Schindelin et al***., 2012). For assessment of patches, cells and patches were segmented using pixel classification in ilastik (***Berg et al***., 2019) to generate two images, one with a mask of the whole cell and one with a mask of the patches, the patch mask also included rings as it wasn’t possible to fully remove rings by means of our training classifier. A custom FIJI script was then used to obtain the area fraction of patches for each segmented cell. Cells that had rings were manually removed from the analysis. Statistics were computed and plotted using R in Rstudio (***Team***, 2022, 2018).

### Western blotting

The four strains were grown overnight in YEA broth. 33 ml of 0.3 OD cells were then lysed using glass beads and vortexing for 1 min and stored on ice and extracted with 500 μl of PBS, 300 μl of this solution was added to 100 μl of 4x Laemelli buffer and was heated at 95°C for 5 mins. They were then loaded and run on 10% SDS-PAGE gels followed by western blotting using the BIO-RAD turbo-blot using the preset mini gel setting. The blot was then blocked with 10% skimmed milk powder in TBS-tween, followed by primary anti-actin antibody in blocking overnight at 4 °C. (MERCK: MAB1501 at 1:1000). The blots were washed 4 times with TBS-Tween for 10 mins, followed by anti-mouse HRP (Cell signaling 7076S at 1:3000) for 1 hour at room temperature.

### Mass spectroscopy

#### Sample preparation

The strains were prepared for SDS-PAGE as above for western blotting. The SDS-PAGE gel was stained with coomassie blue and bands between 35 and 50 kD were cut. The samples were reduced using TCEP and alkylated. They were then digested overnight using Trypsin and concentrated using a speed-vac to 20 μl.

#### NanoLC-ESI-MS/MS Analysis

Reversed-phase chromatography was used to separate tryptic peptides prior to mass spectrometric analysis. Two columns were utilised, an Acclaim PepMap μ-precolumn cartridge 300 μm i.d. x 5 mm 5 μm 100 Å and an Acclaim PepMap RSLC 75 μm x 50 cm 2 μm 100 Å (Thermo Scientific). The columns were installed on an Ultimate 3000 RSLCnano system (Thermo Fisher Scientific). Mobile phase buffer A was composed of 0.1% formic acid in water and mobile phase B 0.1% formic acid in acetonitrile. Samples were loaded onto the μ-precolumn equilibrated in 2% aqueous acetonitrile containing 0.1% trifluoroacetic acid for 5 min at 10 μL min-1 after which peptides were eluted onto the analytical column at 250 nL min-1 by increasing the mobile phase B concentration from 4% B to 25% over 36 min, then to 35% B over 10 min, and to 90% B over 3 min, followed by a 10 min re-equilibration at 4% B.

Eluting peptides were converted to gas-phase ions by means of electrospray ionization and analysed on a Thermo Orbitrap Fusion (Q-OT-qIT, Thermo Scientific) (***Hu et al***., 2005). Survey scans of peptide precursors from 375 to 1575 m/z were performed at 120K resolution (at 200 m/z) with a 50% normalized AGC target and the max injection time was 150 ms. Tandem MS was performed by isolation at 1.2 Th using the quadrupole, HCD fragmentation with normalized collision energy of 33, and rapid scan MS analysis in the ion trap. The MS2 was set to 50% normalized AGC target and the max injection time was 200 ms. Precursors with charge state 2–6 were selected and sampled for MS2. The dynamic exclusion duration was set to 45 s with a 10 ppm tolerance around the selected precursor and its isotopes. Monoisotopic precursor selection was turned on. The instrument was run in top speed mode with 2 s cycles.

#### Data analysis

The data was then analysed using Mascot (***Perkins et al***., 1999) using the following parameters (Enzyme: Trypsin; Fixed modifications: Carbamidomethyl (C); Variable modifications: Oxidation (M), Acetyl (N-term; Mass values: Monoisotropic; Protein mass: Unrestricted; Peptide mass tolerance: ± 10 ppm; Fragment mass tolerance: ± 0.6 Da; Max missed cleavages: 3).

### Spot dilution assay

The different strains were grown in YEA broth overnight at 30 °C. The following day they were diluted and grown to mid-log phase. The cells were then diluted to 0.1 OD (595 nm), 0.01, 0.001, 0.0001 OD on YEA agar plates and incubated at 30 °C for 3 days. They were then scanned on an EPSON perfection V700 Photo scanner.

### Actin Purification

Actin was purified as previously described (***Hatano et al***., 2020). Briefly, MBY12817 cells were grown in MGY liquid medium composed of 1.34% yeast nitrogen base without amino acids (SIGMA: Y0626), 0.4 mg/L biotin and 1% glycerol at 30 °C, 220 rpm to an OD600 of 1.5. They were then pelleted by centrifugation and washed with sterile water. They were then cultured in MM medium containing 1.34% yeast nitrogen base without amino acids (SIGMA: Y0626), 0.4 mg/L biotin and 0.5% methanol at 30 °C, 220 rpm for 1.5-2 days. The cells were pelleted, washed and a frozen in liquid nitrogen. The frozen cells were lysed using a cryo mill (SPEX® SamplePrep: 6870). The lysate was resuspended in an equal volume of 2x binding buffer composed of 20 mM imidazole (pH 7.4), 20 mM HEPES (pH 7.4), 600 mM NaCl, 4 mM MgCl_2_, 2 mM ATP (pH 7.0), 2x concentration of protease inhibitor cocktail (cOmplete, EDTA free : 05056489001, Roche), 1 mM phenylmethylsulfonyl fluoride (PMSF) and 7 mM beta-mercaptoethanol. The lysate was sonicated (5 second with 60% amplitude, QSONICA SONICATORS). The lysate was then centrifuged at 25658g for 5 mins to remove cell debris and a further 1 hour to remove the insoluble fraction. The lysate was then clarified using a 0.22 μm filter. It was then incubated with Ni-NTA beads (Thermo SCIENTIFIC : 88222) at 4 °C for 1 hour. The resin was pelleted and washed repeatedly with 1x Binding buffer and then G-buffer containing 5 mM HEPES (pH 7.4), 0.2 mM CaCl2, 0.01 w/v% NaN3, 0.2 mM ATP (pH 7.0) and 0.5 mM dithiothreitol (DTT). Actin was then cleaved off the beads overnight at 4 °C using 5 μg/ml TLCK treated chymotrypsin (SIGMA: C3142-25MG) which was then inactivated by 1mM PMSF. The beads were pelleted and the supernatant was concentrated using a 30kDa cutoff membrane to 0.9 ml. The actin was polymerized using 100 μl 10x MKE solution composed of 20 mM MgCl_2_, 50 mM ethylene glycol tetraacetic acid (EGTA) and 1 M KCl for 1 hour at room temperature and pelleted down by ultracentrifugation at room temperature (45000 rpm for 1 hour, Beckman TLA-55 rotor). The pellet was then resuspended in 1x G-buffer and dialyzed against 2L of G-buffer over 48 hours at 4 °C.

Actin for EM was polymerized by mixing, 20 μl 20 μM G-actin, 8 μl 10x MKE and 52 μl 5 mM HEPES-KOH pH 7.4 containing 0.2 mM ATP and 0.5 mM DTT and incubating at RT for 1 hour.

### Myosin purification

Porcine cardiac myosin for motility assays was purified as described previously (***Murakami et al***., 1976). The protocol was not modified. The myosin was stored in 50% glycerol at -20 °C.

### Actin activated myosin ATPase assay

Actin activated myosin ATPase activity assay was performed using EnzChek ATPase kit (ThermoFisher). The assay was performed in 100 μl in a cuvette. Reaction buffer (5mM KCl, 0.6 mM MgCl_2_, 10 mM PIPES) was mixed with 2-amino-6-mercapto-7-methylpurine riboside (MESG), purine nucleoside phosphorylase (PNP), polymerized actin, bovine cardiac myosin S1 (25 μg/ml) and ATP (40mM). As soon as actin, myosin and ATP was mixed with other components absorbance of the assay mixture was measured at 360 nm for 15 minutes. Then rate of absorbance change was calculated for all conditions.

### Motility assay

#### Sample preparation and imaging

The movement of actin filaments on myosin was assessed using a motility assay. Acetylated actin or arginylated actin were co-polymerized with 5% fluorescently labelled Ac-actin (Alexa Fluor 488 (Invitrogen)). While R-β-actin was labelled with Alexa Fluor 488 (Invitrogen). Actin was polymerized at room temperature for 1 hour by adding MEK polymerization buffer (20 mM MgCl_2_, 50mM glycol-bis(2-aminoethylether)-N,N,N′,N′-tetraacetic (EGTA) and 1 M KCl. Glass coverslips were cleaned in 2% Helmanex II for 60 mins at 60 °C and then in 3 M NaOH for 30 min at 60 °C. Both steps were performed in an ultrasonic bath. Cleaned coverslips were rinsed with pure water and air dried. Then coverslips were coated with silicone by immersing them in 5% (v/v) Sigmacote (Sigma-Aldrich) in heptane. The coverslips attached to microscopy slides using three stripes of a double-sided cellotape. Each coverslip had two 2 mm wide channels. Porcine cardiac myosin (1 mg/ml) was loaded into both channels and incubated for 10 minutes at room temperature. Then the channels were washed with 20 μl running buffer (25 mM KCl, 4 mM MgCl_2_, 1 mM ATP, 10 mM DTT, 25 mM Imidazole pH 7.4, 1 mg/ml BSA) twice. The channels were incubated with running buffer for 2 minutes. 20 μl of running buffer supplemented with 20-40 nM of polymerized actin were loaded into channels. First channel was loaded with acetylated actin while the second was with arginylated-β-actin. The channels were sealed with nail polish. The samples were imaged with a confocal fluorescence microscope. Images were taken every 100 msec for 20 sec. Actin filament movements were analyzed with Fiji and Trackmate plugin (***Ershov et al***., 2022).

#### Image analysis

The pipeline for image analysis of detachment and wobbling of actin filaments during motility is illustrated in Fig. S2. E. To visualize detachment and wobbling, Gaussian blurring with a radius of 2 pixels was first applied to the images. The ratios between the pixel intensities of the n-th and (n+1)-th images of the movie were then calculated. These ratio images were maximum-projected through the time points and presented on a linear scale from 1.3 to 3 using the pseudo-color ‘FIRE’ lookup table. For scoring the detachment and wobbling events, the actin filaments and background in the original movie were first segmented using Trainable Weka Segmentation (***Arganda-Carreras et al***., 2017) in Fiji. The mean and standard deviation of the background pixel intensities were measured. Spots in the ratio images that exceeded a threshold defined as

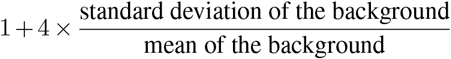

were counted as detachment or wobbling events.

### Electron Microscopy

Samples were prepared by applying the polymerized R-β-Actin actin to freshly glow-discharged Quantifoil 3.5/1 on 200 mesh copper grids (Quantifoil GmbH, Germany), using a Leica GP2 (Leica Microsystems GmbH, Germany). Grids were screening on a JEOL2200FS with K2 camera. Data collection was carried out on the ThermoFisher Scientific Titan Krios using the Gatan K2 Direct Electron Detector, operated at a nominal magnification of 75000x, resulting in a calibrated pixel size of 1.08 Å/pix. Micrographs were recorded as movies comprising 39 individual frames recorded over 60 seconds and with a dose rate of 0.7 electrons per pixel per second, giving a total dose per image of 42 electrons/pixel.

### Image processing and model building

All data processing was done using RELION3.0.9 (***Zivanov et al***., 2018). Image stacks were motion-corrected and summed using MotionCor2 (***Zheng et al***., 2017) and CTF parameters were calculated using Gctf (***Zhang***, 2016). Initially, filament segments were manually selected from a subset of micrographs and 2D classes were calculated. These were then used to optimise the settings for autopicking on a subset of micrographs with a range of defocus values, before running autopicking on all images. Bad particles were removed using 2D and 3D classification, before refining the helical parameters in 3D refinement. Resolution was estimated using the Fourier shell correlation 0.143 criterion and local resolution maps were produced using LocResMap.

PDB model 3J8I (Galkin et al 2015) was fitted into the density map to assess similarity to previous structures. Model building was performed in COOT (***Emsley and Cowtan***, 2004), mutating the primary sequence to account for the difference between chicken actin and human actin. Refinements were carried out in reciprocal space using REFMAC (***Murshudov et al***., 1997) and geometry was analysed using Coot and Molprobity (***Chen et al***., 2010).

### Data deposition

The map for R-β-actin has been deposited in the electron microscopy data bank – accession number EMD-16776 and in the protein data bank with the PDB ID 8COG.

### A model of the actomyosin complex

An actin-myosin 1:1 complex was modelled based on chains A and F of a cryo-EM structure (PDB:5JLH (***von der Ecken et al***., 2016)). The missing loops of non-muscle myosin IIC were modelled with MODELLER (***Šali and Blundell***, 1993) via UCSF Chimera (***Pettersen et al***., 2004). Molecular dynamic simulation was performed with GROMACS (https://www.gromacs.org/ (Páll, S.)(***Páll et al***., 2015)) using the output from CHARMM-GUI (https://www.charmm-gui.org/ ***Jo et al***. (2008); ***Lee et al***. (2016)) with N-terminal acetylation of the gamma actin. Coulomic surface colouring was done by UCSF Chimera.

## Acknowledgments

We acknowledge the University of Warwick Advanced Bioimaging Research Technology Platform supported by BBSRC ALERT14 award BB/M01228X/1 and the Midlands Regional Cryo-EM Facility, hosted at the Leicester Institute for Structural and Cellular Biology, for use of the Titan Krios, supported by MRC award reference MC_PC_17136. We thank TJ Ragan for his assistance with data collection. We thank the University of Warwick, School of Life Science, Proteomics Core facility and Cleidiane Zampronio for processing the mass spectrometry samples. The authors have no additional competing financial interests. We also thank the Henriques Lab and eLIFE for their Latex templates (license CC. BY. 4.0) that we modified and used on Overleaf to generate this manuscript. This work was supported by a Wellcome Trust Senior Investigator Award (WT101885MA; MKB), a Wellcome Trust Collaborative Award in Science (203276/Z/16/Z; MKB), a European Research Council Advanced Grant (ERC-2014-ADG No. 671083; MKB), and a collaborative BBSRC award (BB/S003789/1; MKB, MM, KS).

## Description of Videos

**Video 1**. Time-lapse movies of motility assays for the movement of F-actin. Fluorescent filamentous Ac-β-Actin or R-β-Actin (equal amounts) were loaded on immobilized porcine cardiac myosin. Movement of actin filaments was demonstrated using MTrackJ plugin in Fiji. Scale bars are 10 μm.

**Video 2**. Time-lapse movies of cells from the MEEE and REE strains in the rlc1-GFP background as indicated showing mCherry-Atb2 (magenta) and Rlc1-3GFP (green). Images were captured at 3 min intervals. Scale bar is 1 μm. Montages of the individual movies are shown in Figure 4 A.

## Supplementary Figures

**Fig. S 1.**
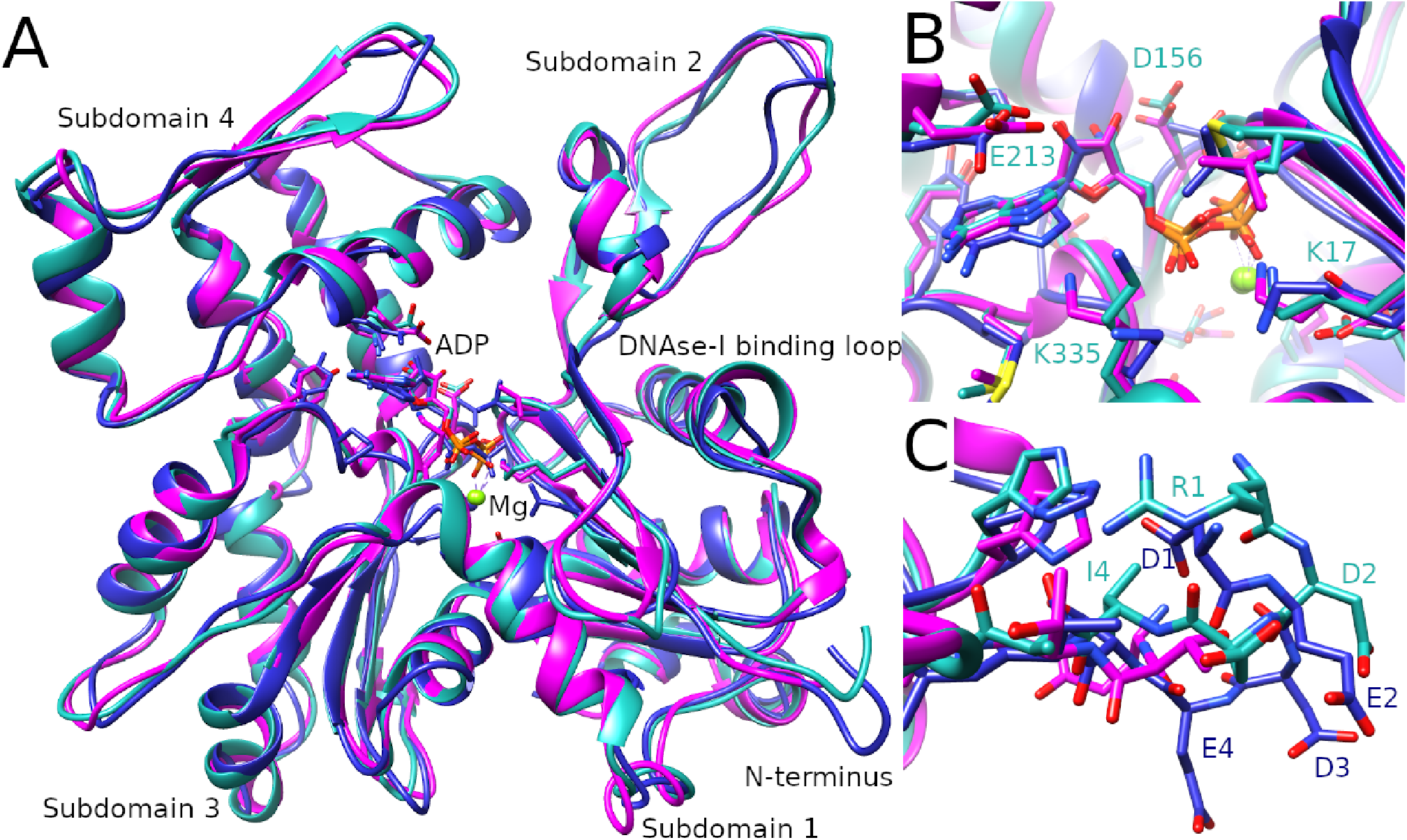
Comparison of the R-β-actin, chicken and rabbit skeletal muscle actin structures. (A) Ribbon diagram comparing the structure of R-β-actin presented here (cyan) with previously published structures of chicken (navy) and rabbit (magenta) skeletal muscle actin. (B) Close up view comparing the structure of the active site of R-β-actin with previously published structures of chicken and rabbit skeletal muscle actin. (C) Close up view comparing the N-terminal sections of R-β-actin to that of chicken and rabbit skeletal muscle actin.

**Fig. S 2.**
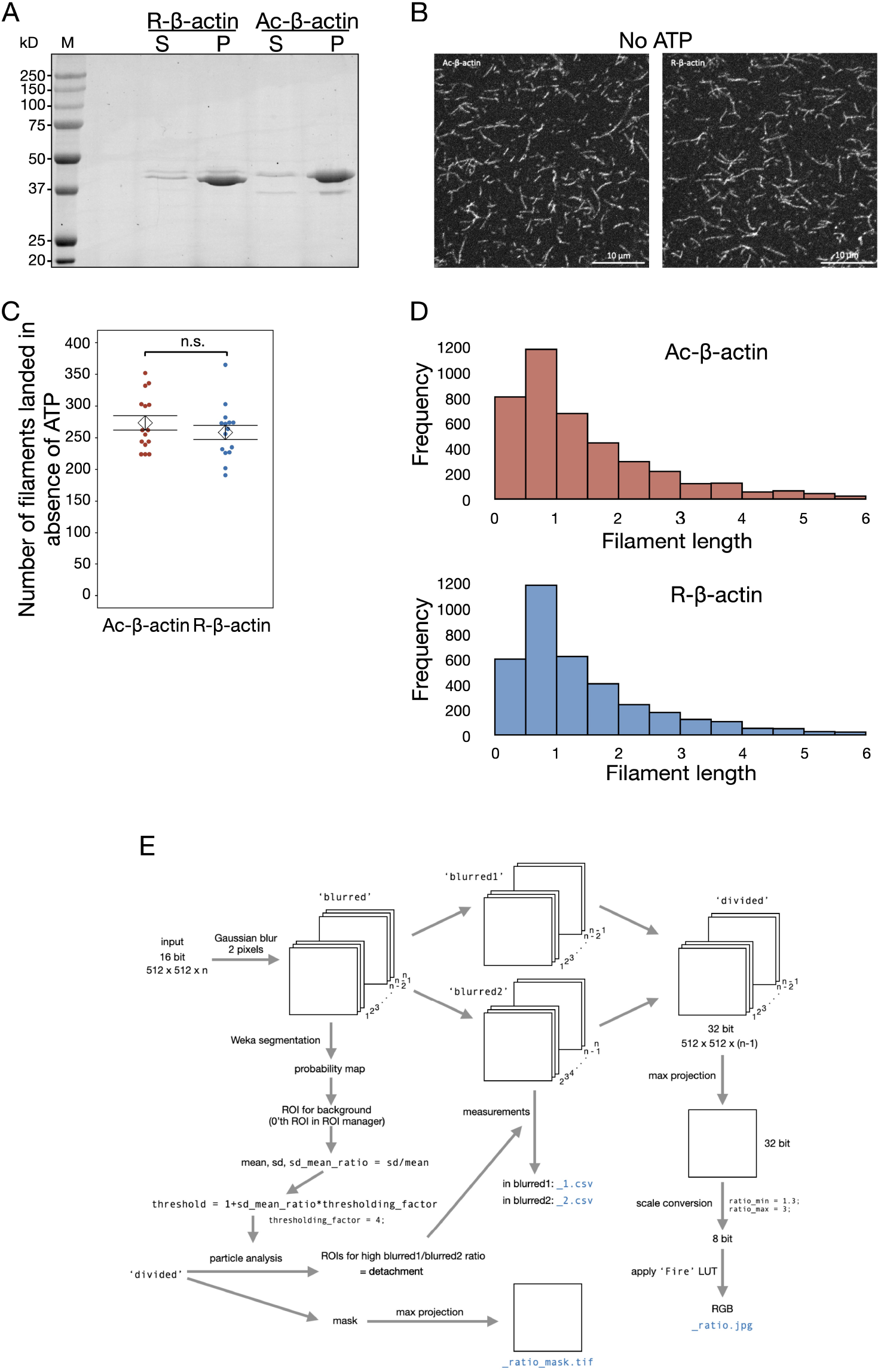
The degree of assembly into filaments and the length distribution of the filaments arginylation of actin. (A) The purified Ac- and R-actin were polymerized in vitro and ultracentrifuged at 100,000 xg for 1 h. The proteins in the supernatent (S) and pellet (P) were assed by SDS-PAGE and coomassie staining. (B) Alexa Fluor-488 -labeled Ac- and R-actin filaments were loaded on the surfaced coated with cardiac myosin in the absence of ATP. (C) The number of the landed filaments. (D) The length distributions of the landed filaments. (E) Schematic of the image anlaysis pipeline to detect and analyze detachement/ wobbling of actin filaments in the surface gliding assay. The pixels that showed a large difference from the previous time frame were detected by dividing the (i+1)-th images with the i-th images after smoothing with gaussian blurring. The maximum projections through the time frames are shown in a pseudocolor as shown in Fig. 3. D. For quantification of the detachment/wobbling events, a threshold was calculated based on the mean and the standard diviation of the background areas in each movie and the spots above this threshold value in the (i+1)-th/i-th ratio movie were detected.

**Fig. S 3.**
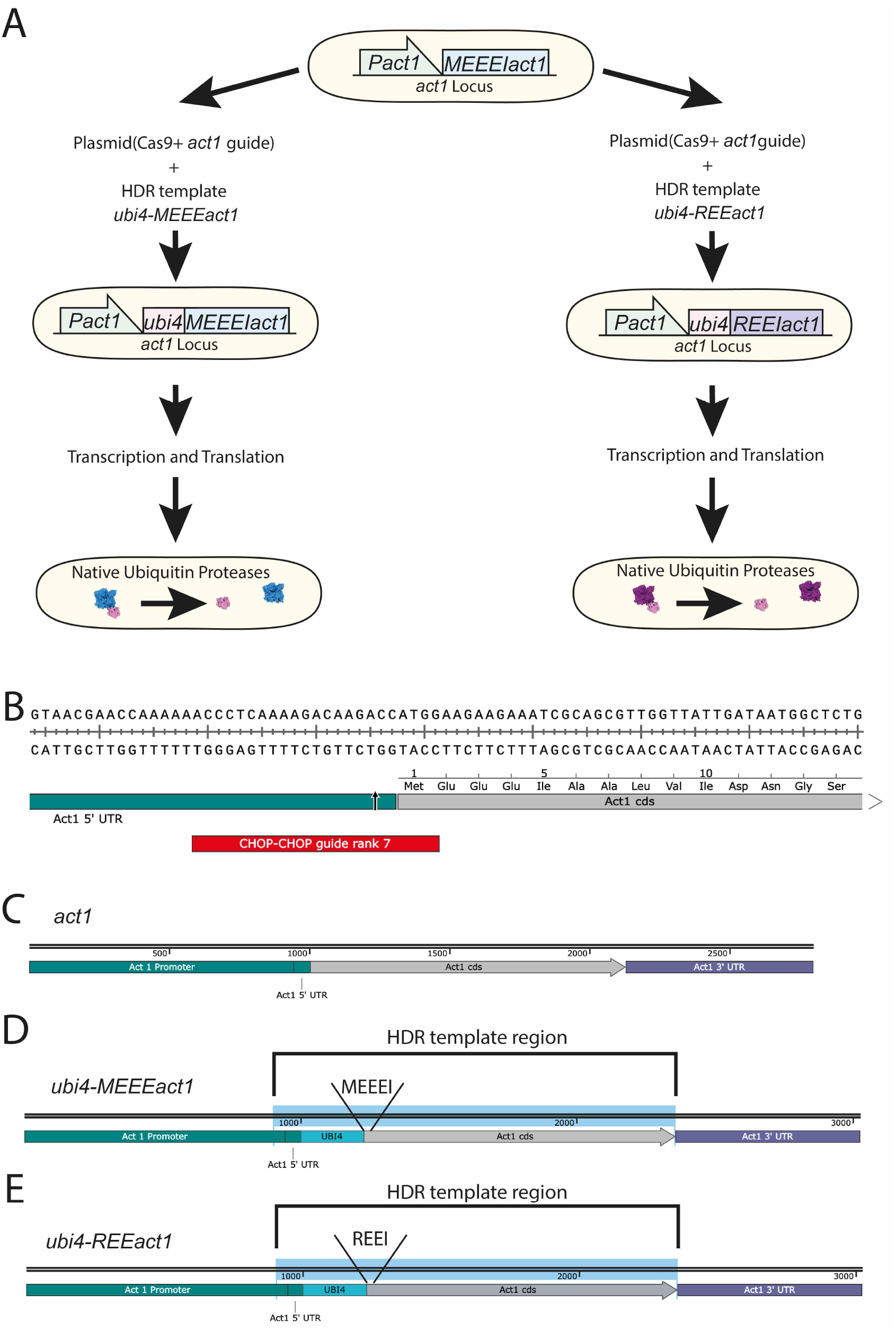
Approach used to generate the R-Actin and the control *S.pombe* strains. (A) Schematic of the approach used to generate the REE and MEEE strains. (B) Sequence of the junction between the *act1* promoter and the *act1* coding sequence (cds) showing the location of the chosen guide RNA target (red) designed in CHOP-CHOP and the expected Cas9 cut site (arrow). (C) Schematic of the *act1* locus. (D) Schematic of the expected *ubi4-MEEEact1* incorporation at the native *act1* locus including the HDR template region that was used to generate the strains. (E) Schematic of the expected *ubi4-REEact1* incorporation at the native *act1* locus including the HDR template region that was used to generate the strains.

**Fig. S 4.**
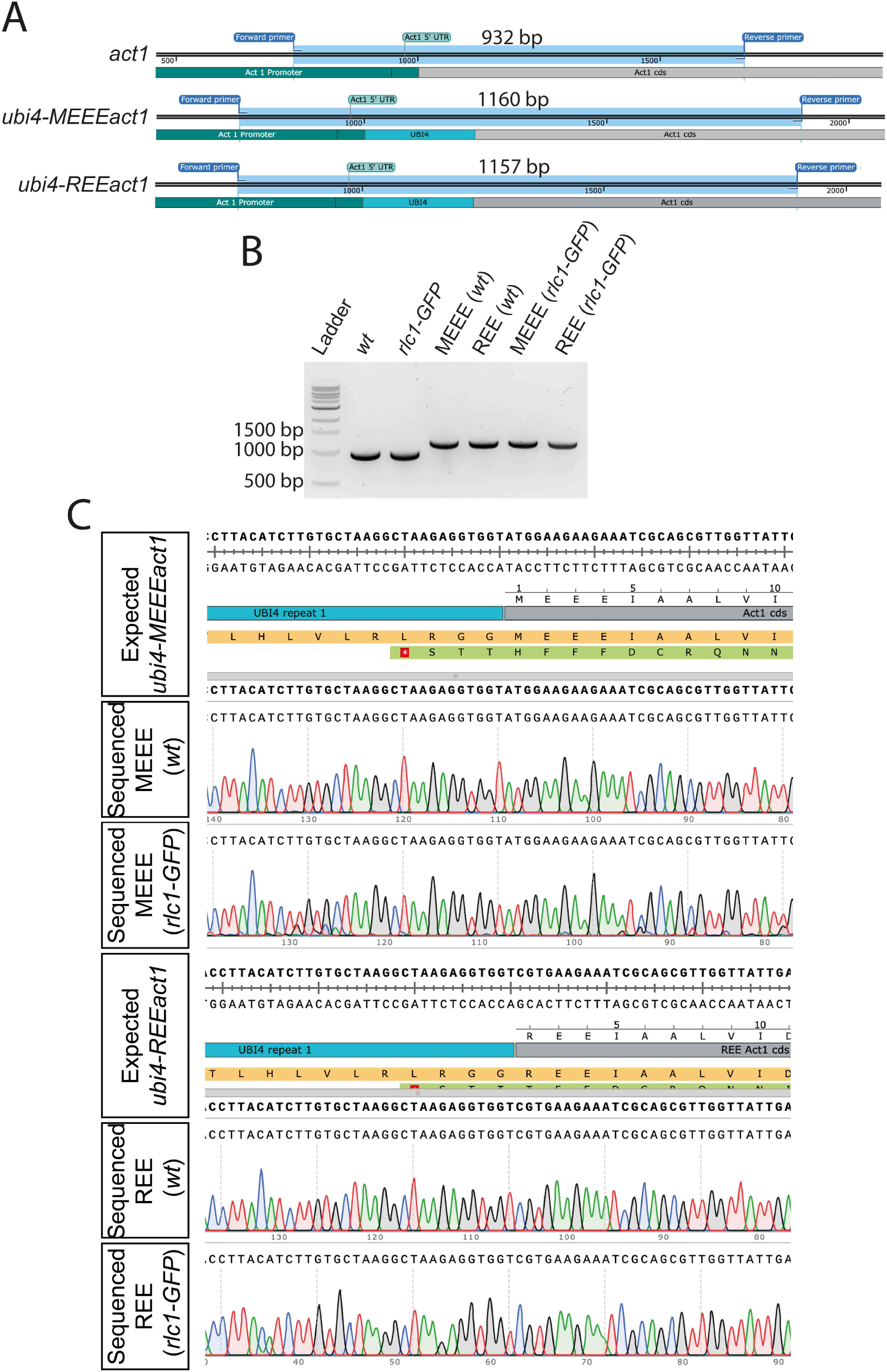
Validation of the R-Actin and control *S.pombe* strains. (A) Schematic of the pcr products and their sizes for the *act1, ubi4-MEEEact1* and *ubi-REEact1* strains using the same pair of primers for genotyping. (B) 1% Agarose gel image of the pcr products of the different strains using the genotyping primers. Note the presence of a single band of the expected sizes for all the strains. (C) Sequence alignment results for the 4 *ubi4* strains as compared to their respective expected sequences as derived from Snapgene® software.

**Fig. S 5.**
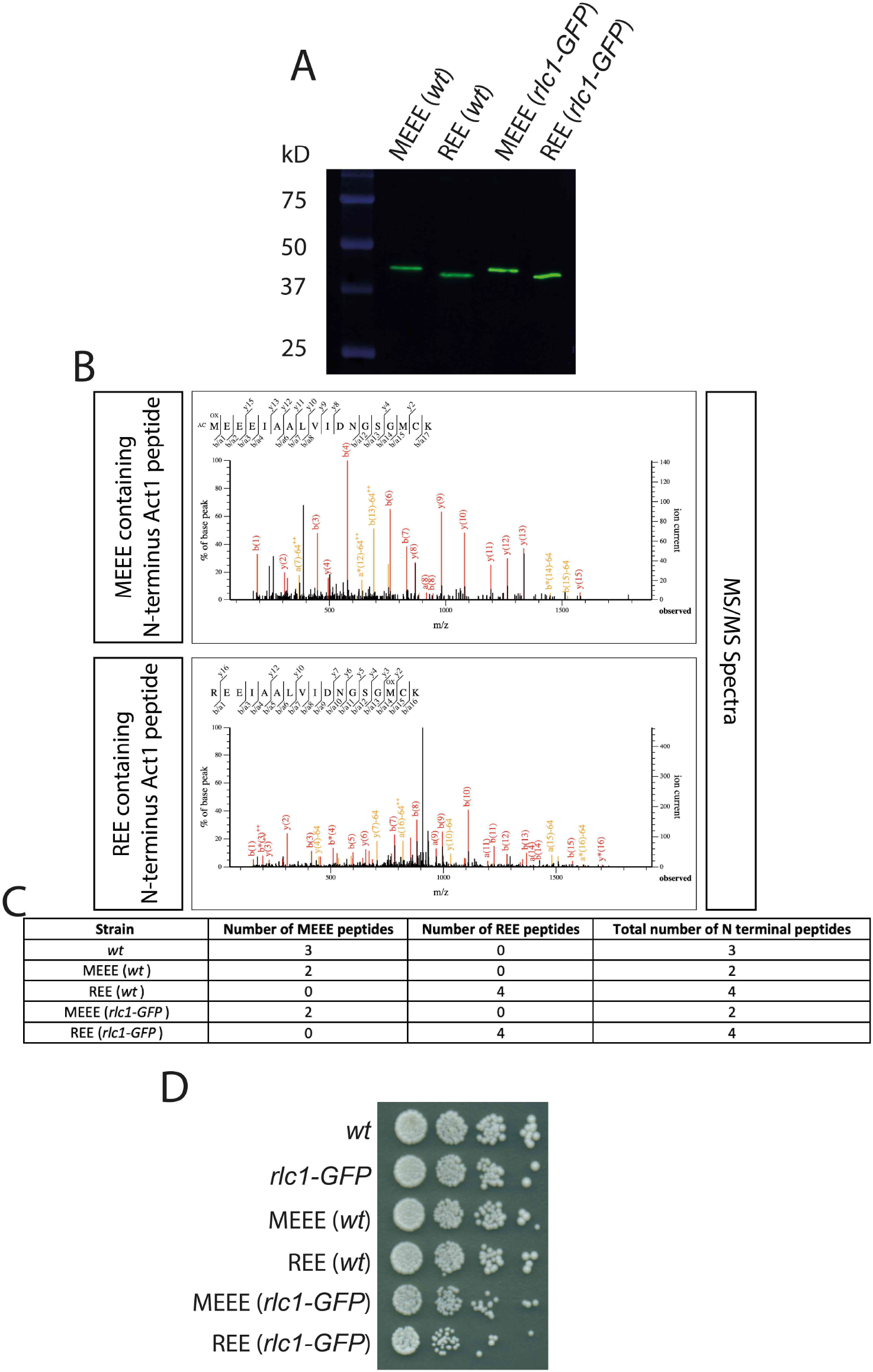
Proteomic analysis of actin and spot assays of the R-actin and control *S.pombe* strains. (A) Western blot using an anti-actin antibody for the 4 *ubi4* strains. The actin bands are at the expected size of 42 kD of the actin product post Ubi4 tag cleavage. Also note the differences in migration of actin between the MEEE and REE strains (B) MS/MS spectrum for an MEEE peptide from the MEEE (wt) strain and for an REE peptide from the REE (wt) strain. (C) Table of the number of MEEE or REE peptides observed for the different strains. (D) Spot assay showing the growth of 10-fold serial dilutions of the indicated strains that were spotted on YEA plates and grown at 30°C for 3 days.

**Fig. S 6.**
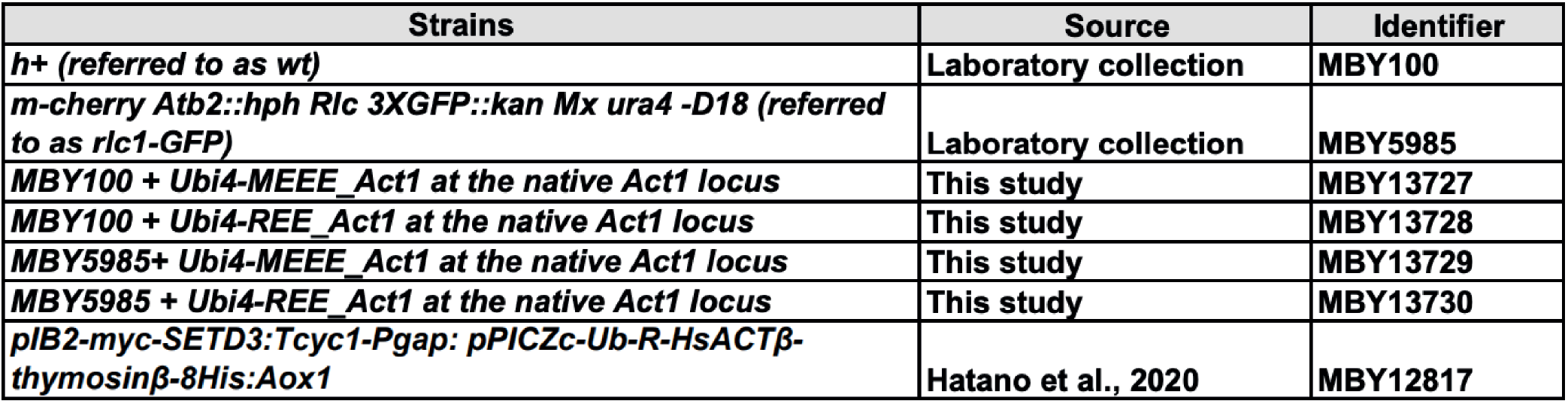
List of *S.pombe/P.pastoris* strains used in this study.

